# An oxide transport chain essential for balanced insulin signaling

**DOI:** 10.1101/430637

**Authors:** Xiangdong Wu, Keyang Chen, Kevin Jon Williams

## Abstract

Patients with overnutrition, obesity, the atherometabolic syndrome, and type 2 diabetes mellitus exhibit *imbalanced insulin action*, also called pathway-selective insulin resistance. To control glycemia, they require hyperinsulinemia that then overdrives ERK and hepatic de-novo lipogenesis. We recently reported that NADPH oxidase-4 regulates balanced insulin action. Here, we show that NADPH oxidase-4 is part of a new limb of insulin signaling that we abbreviate “NSAPP” after its five major proteins. The NSAPP pathway is an oxide transport chain that begins when insulin stimulates NADPH oxidase-4 to generate 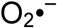. NADPH oxidase-4 hands 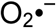 to superoxide dismutase-3 for conversion into H_2_O_2_. The pathway ends when aquaporin-3 channels H_2_O_2_ across the membrane to inactivate PTEN. Disruption of any component of the NSAPP chain, from NADPH oxidase-4 up to PTEN, leaves PTEN persistently active, thereby producing the same deadly pattern of imbalanced insulin action seen clinically. Unraveling the molecular basis for NSAPP dysfunction in overnutrition has now become a top priority.

## Text

Eighty-seven years ago, Falta & Boller published that human patients with what we now call type 2 diabetes mellitus (T2DM) exhibit a characteristic resistance to the glucose-lowering actions of insulin.^1–3^ Just over two decades ago, however, Hellerstein et al. reported that administration of insulin to patients with T2DM still vigorously activates hepatic de-novo lipogenesis, despite concurrent resistance to the ability of insulin to suppress hepatic glucose production.^4^

Thus, the real clinical issue in T2DM and related syndromes of overnutrition is *imbalanced insulin action*, which is also known as pathway-selective Insulin <u>r</u>esistance and <u>r</u>esponsiveness (SEIRR).^3–7^ Individuals with SEIRR require compensatory hyperinsulinemia to control plasma glucose concentrations. The result is overdrive of those pathways that remain insulin-responsive, such as ERK activation^5^ and hepatic de-novo lipogenesis.^4,8^ The effects are easily summarized: if hyperinsulinemia does something undesirable in a tissue or organ (red in our schematic Fig. 1), that effect remains responsive in T2DM and other syndromes of overnutrition. If hyperinsulinemia might do something beneficial (blue in Fig. 1), that effect becomes insulin-resistant. From the standpoint of human health, it is the worst possible combination of effects.^3,6,7^ Many studies by now have demonstrated this harmful pattern of imbalanced insulin action in obese non-diabetic and T2DM humans^4,5,8^ and in several spontaneously hyperphagic rodent models on normal chow.^6,9–15^ Nevertheless, the molecular basis for imbalanced insulin action in states of overnutrition has remained a major unknown in metabolic research.

We recently discovered that healthy, balanced insulin signaling requires transient activation of the NADPH oxidase-4 (NOX4), generation of H_2_O_2_, and hence inactivation of PTEN and other redox-sensitive members of the protein tyrosine phosphatase (PTPase) gene family (encircled pathway in Fig. 1).^6,7^ Hyperphagia and obesity in animals disrupts NOX4 signaling in liver in response to insulin, resulting in persistently active hepatic PTEN that blocks normal signaling to AKT.^6^ Unexpectedly, despite the abundance of enzymatically active PTEN in livers of hyperphagic obese animals, we found that insulin still stimulated full two-site phosphorylation (p) of ERK (pT202/pY204-ERK); hepatic generation of an unusual monophosphorylated form of AKT at Thr308 (pT308-AKT) with only weak phosphorylation at Ser473; robust pT308-AKT-mediated phosphorylations of PRAS40 and GSK3ß (upstream of de-novo lipogenesis in Fig. 1); but impaired AKT-mediated phosphorylation of FOXO1 (required for suppression of hepatic gluconeogenesis, Fig. 1).^6^ Monophosphorylated pT308-AKT is enzymatically active on many immediately downstream substrates, but unless AKT has undergone two-site phosphorylation (pT308/pS473-AKT), it cannot act on FOXO1.^3,6,7,15,16^

**Figure 1:**
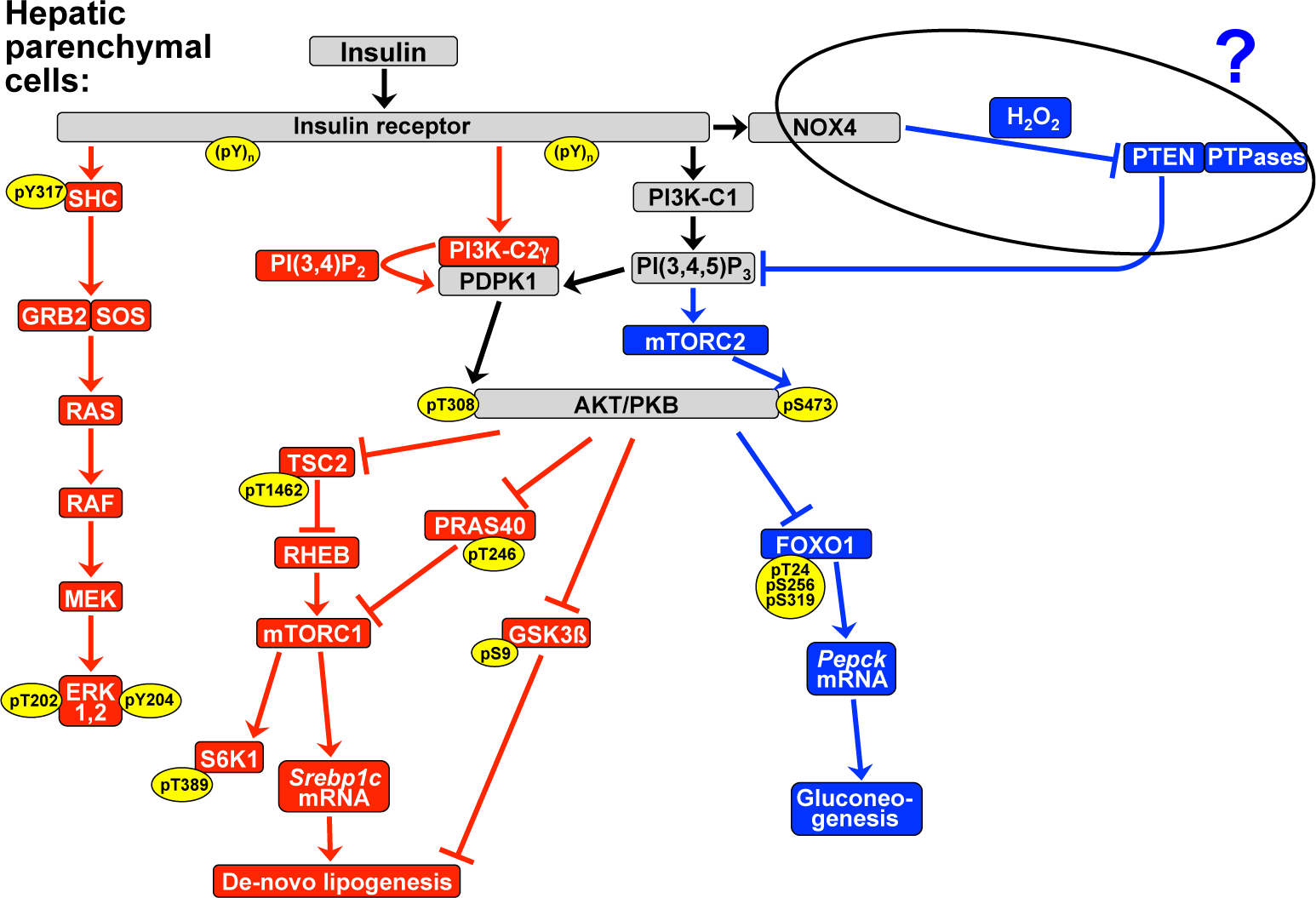
Prior model of the key limbs of insulin signaling in hepatic parenchymal cells. Pathways shown in **red** can cause side-effects characteristic of overnutrition, obesity, the atherometabolic syndrome, and T2DM. We administer exogenous insulin and insulin secretagogues to simulate the **blue** pathways. **Pointed arrowheads** indicate stimulation of the immediately downstream molecule or process; **flat arrowheads** indicate inhibition. Any chemical compound written next to the shaft of an arrow transmits the signal. Yellow ovals indicate sites that become phosphorylated (p) upon insulin stimulation. AKT/PKB, protein kinase B; mTORC, mammalian target of rapamycin complex; PI, phosphatidylinositol; PI3K, PI 3’-kinase; PI3K-C1, class I PI3Ks; PI3K-C2γ, PI3K class II, gamma. Other protein and mRNA abbreviations follow OMIM and HGNC. The encircled pathway from NOX4 to PTEN is the focus of the current study. Adapted from references^3,6^ with permission.

To investigate causal relationships, we showed that artificial impairment of NOX4 in cultured liver cells recapitulates all features of imbalanced hepatic insulin signaling that we documented in vivo.^6^ Thus, the defect in insulin-stimulated activation of the NOX4 pathway that we demonstrated in livers of hyperphagic obese animals^6^ may finally explain Hellerstein et al.’s pioneering finding from over 20 years ago^4^ that the administration of insulin to human patients with T2DM still triggers hepatic de-novo lipogenesis, but without controlling hepatic glucose metabolism.

Despite its explanatory power, however, our model of the NOX4 pathway displayed schematically in Fig. 1 leaves two mysteries. First, the catalytic site of NADPH oxidases generates superoxide 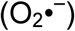, yet NOX4 appears to produce H_2_O_2_, ^17,18^ a molecule with considerably different physical properties. Second, NADPH oxidases emit their product away from the cytosol,^19^ but PTEN and PTPases, the key targets of NOX4 in this context, reside within the cytosol. In the current study, we focused on these two mysteries. This work was presented at the American Diabetes Association Scientific Sessions,^20-22^ and the full initial manuscript was uploaded onto the bioRXiv (28 September 2018).

### Oxide hand-off from NOX4 to SOD3

The finding that NOX4 generates H_2_O_2_ and not superoxide has resisted biochemical explanation.^17,18^ To provide a model that incorporates prior findings from enzymology and insulin signaling,^3,6^ we had hypothesized that a nearby superoxide dismutase (SOD) must be required to promptly convert 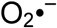 from NOX4 into longer-lasting H_2_O_2_ that can then physically reach PTEN to inactivate it.^7^ The mammalian genome contains exactly three SODs: the cytosolic SOD1, the mitochondrial SOD2, and the secreted SOD3 (also known as the extracellular SOD, ecSOD).^23^ Thus, our initial approach was to survey all three SODs using co-immunoprecipitations (Fig. 2). We found that NOX4 in McArdle hepatocytes does not detectably co-immunoprecipitate with either the cytosolic SOD1 or the mitochondrial SOD2 (Fig. 2a). These results are consistent with a prior report indicating that NOX4 does not associate with intracellular SODs, although that study did not examine the extracellular SOD3.^18^

**Figure 2:**
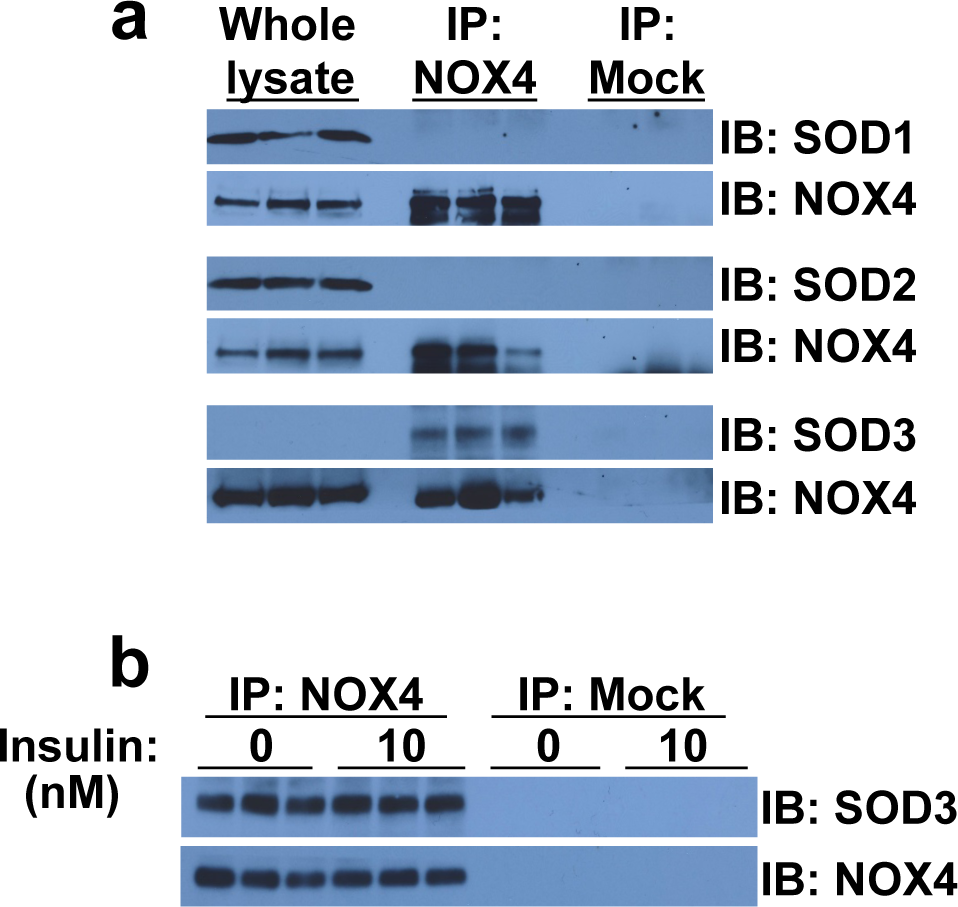
NOX4 and SOD3 form a novel complex together. **a**, Immunoprecipitation (IP) of NOX4 from homogenates of McArdle hepatocytes, followed by immunoblots (IB) for SOD1, SOD2, and SOD3. To verify immunoprecipitation and loading, the same membranes were stripped and reprobed to detect NOX4 (IB: NOX4). To confirm that the antibodies against SOD1 and SOD2 detect their targets in hepatocytes, whole-cell homogenates before immunoprecipitation were included on the immunoblots. Also included are mock immunoprecipitations, performed on cell homogenates but without the anti-NOX4 antibody. The band for SOD3 in whole-cell homogenates becomes easily visible with longer exposure. **b**, Coimmunoprecipitation of NOX4 and SOD3 from homogenates of McArdle hepatocytes that we obtained after exposing the cells to 0 or 10 nM insulin for 10 minutes.

In contrast, we found that NOX4 robustly co-immunoprecipitates with SOD3 in McArdle hepatocytes (Fig. 2a). In these experiments, both NOX4 and SOD3 were endogenous, i.e., at normal levels of expression. The SOD3 is an extracellular and cell-surface enzyme that had no previously known relationship to NOX4.^3,23,24^ Thus, NOX4 and SOD3 form a novel complex together.

Because insulin^25^ and insulin-like growth factor-1 (reference^26^) were reported to affect the subcellular localization of NOX4 in non-hepatic cell types, we determined if insulin regulates the tight association of NOX4 with SOD3. Fig. 2b shows that the association of NOX4 with SOD3 in McArdle hepatocytes is unaffected by exposure of the cells to insulin.

Most importantly, we found that knockdown of SOD3 in cultured liver cells recapitulates the harmful pattern of imbalanced insulin signaling seen in the liver in overnutrition and in cultured hepatocytes after artificial disruption of NOX4.^6^ In McArdle hepatocytes pre-treated with *Sod3* siRNA to render them deficient in SOD3 protein (Fig. 3a), insulin failed to stimulate an intracellular H_2_O_2_ burst (Fig. 3b) or to inactivate PTEN (Fig. 3c). Prior work had not examined the subcellular distribution of normal, endogenous, hormonally induced H_2_O_2_ (reviewed in reference^3^). Unexpectedly, our confocal images of individual control hepatocytes showed that the insulin-stimulated burst of intracellular H_2_O_2_ was localized mainly to a perinuclear area (Fig. 3b, inset), indicating spatial as well as temporal regulation.^3^ Insulin stimulation of SOD3-deficient hepatocytes provoked robust phosphorylations of ERK at Thr202 and Tyr204, as well as ample production of pT308-AKT, with only weak phosphorylation at the Ser473 site of AKT (Fig. 3d). Fig. 3e displays the effects of SOD3 knock-down on targets immediately downstream of AKT – namely, continued insulin-stimulated phosphorylations of TSC2, PRAS40, and GSK3ß (upstream of hepatic de-novo lipogenesis in Fig. 1), but poor insulin-stimulated phosphorylation of FOXO1 (required for suppression of gluconeogenesis). Thus, without SOD3, all of the red pathways in Fig. 1 remain insulin-responsive, while the blue pathways become insulin-resistant. These results identify SOD3 as essential for normal, balanced insulin signaling.

**Figure 3:**
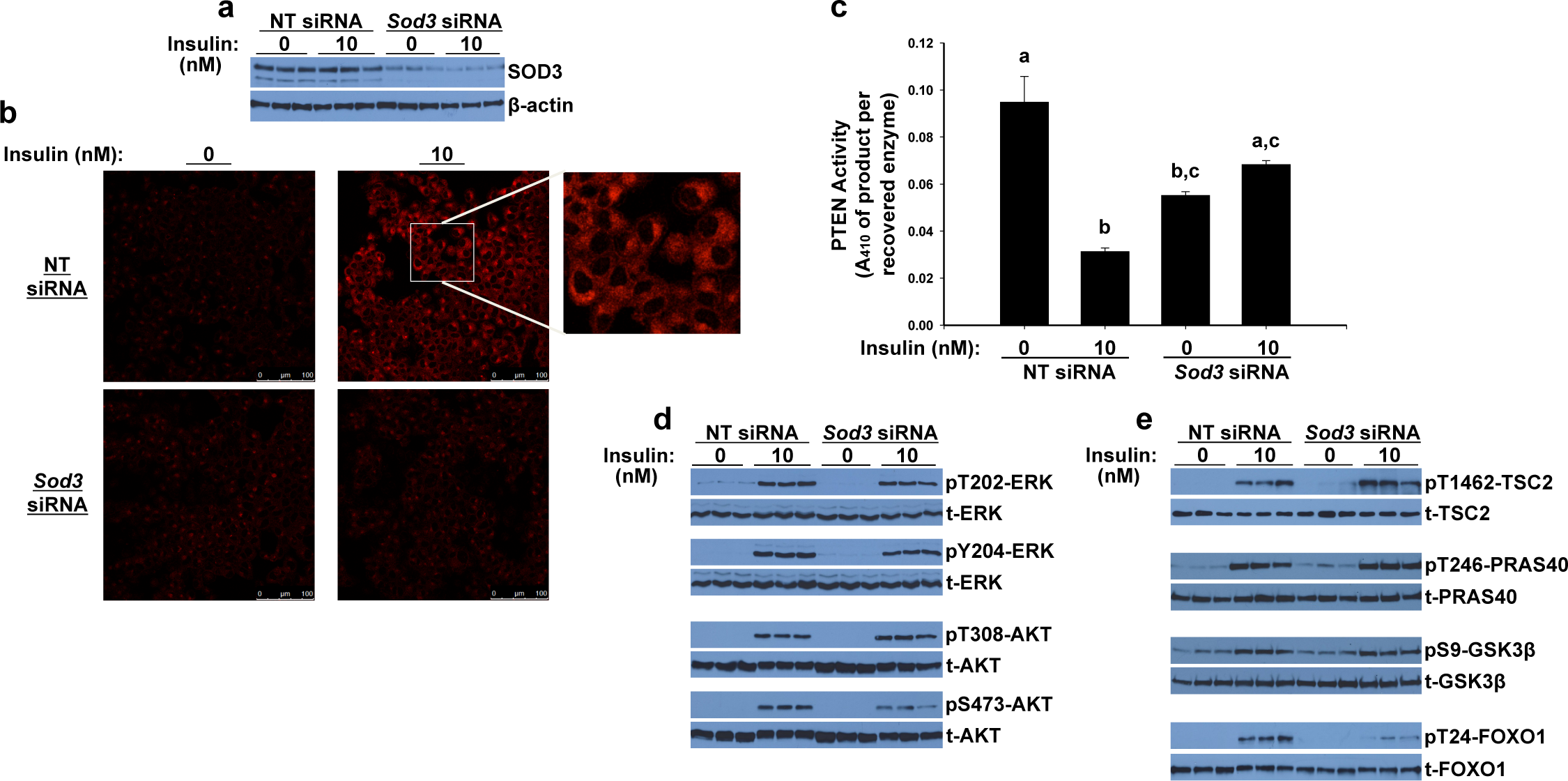
Knock-down of the extracellular SOD3 in cultured McArdle hepatocytes recapitulates the harmful pattern of imbalanced insulin signaling seen in overnutrition or after artificial disruption of NOX4. McArdle hepatocytes were pretreated with nontarget (NT) control or *Sod3* siRNAs, as indicated. Cells were exposed to 0 or 10 nmol/L insulin for 10 minutes, and then harvested. Immunoblots came from a single set of cultured cells. Each lane represents a separate well. **a**, Anti-SOD3 immunoblot. **b**, Insulin-stimulated production of intracellular H_2_O_2_, assessed by confocal images of CellROX Deep Red fluorescence. We definitively identified the intracellular CellROX signal as H_2_O_2_ by its quenching by catalase (not shown). **c**, PTEN activities in cell homogenates, assayed under strictly anaerobic conditions (means±SEMs, n=6, P<0.0001 by ANOVA; columns that do not share a lowercase letter differ by Tukey’s test, P<0.01). **d**, Insulin-stimulated phosphorylations of key sites on ERK and AKT. Total (t-) ERK, AKT are also shown. **e**, Insulin-stimulated phosphorylations of targets directly downstream of AKT, in de-novo lipogenic pathways (pT1462-TSC2, pT246-PRAS40, pS9-GSK3ß) and in glucose-lowering (pT24-FOXO1). Nomenclature follows Fig. 1.

### Molecular basis for NOX4 binding to SOD3

Takac et al.^18^ previously reported that H_2_O_2_ production by NOX4 depends on two short inserts in its non-cytosolic E-loop that are absent from sequence alignments with NOX1 or NOX2. To explain their findings, the authors proposed that the E-loop inserts in NOX4 might possess intrinsic dismutase activity.^18^ Our results in Fig. 3b do not support this proposal, because insulin stimulated no detectable burst of H_2_O_2_ after SOD3 knock-down, yet endogenous NOX4 in those cells still possessed its two E-loop inserts. Thus, we hypothesized an alternative explanation – namely, that the E-loop inserts mediate the tight binding of NOX4 to SOD3 and thereby change the apparent product of NOX4 from 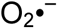 to H_2_O_2_.

**Figure 4:**
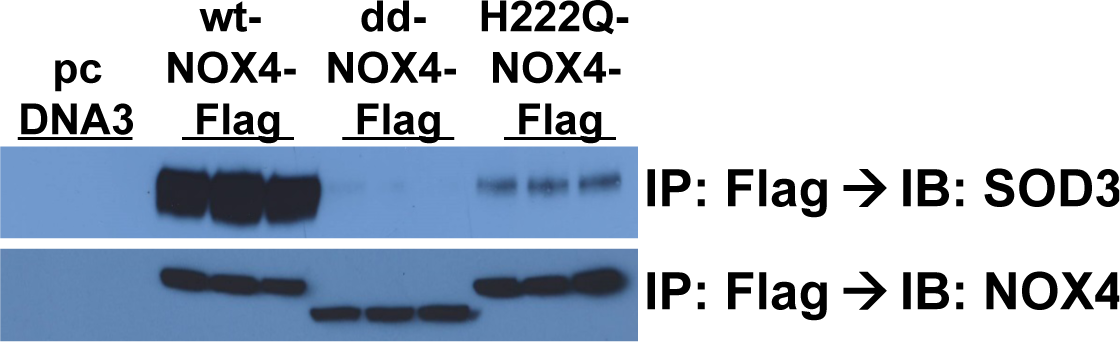
The two inserts in the extracellular E-loop of NOX4, and particularly histidine-222 in the first of those inserts, mediate the normal, robust binding of NOX4 to SOD3. McArdle hepatoma cells were prepared that express, one at a time, an empty pcDNA3.1(-) vector (pcDNA3) or the vector with our Flag-tagged NOX4 constructs: wild-type human NOX4 (wt-NOX4-Flag), a double-deletion mutant that lacks both E-loop inserts (dd-NOX4-Flag), and a site-directed mutant of the key histidine-222 residue to glutamine (H222Q-NOX4-Flag). Shown are immunoprécipitations of the Flag epitope (IP: Flag), followed by immunoblots (IB) to detect endogenous SOD3 protein and immunoreactive NOX4 protein. Each lane represents a separate culture well. As expected, dd-NOX4-Flag appears at a lower molecular weight than do the other NOX4 constructs.

To test our hypothesis, we generated a series of human NOX4 expression constructs, each linked at the 3’ end of the coding region (C-terminus of the resulting protein) to a sequencing encoding the Flag epitope tag. We stably expressed our constructs in McArdle hepatocytes: wild-type human NOX4 (wt-NOX4-Flag), a double-deletion mutant that lacks both E-loop inserts (dd-NOX4-Flag), and a site-directed mutant of the key histidine-222 residue identified by Takac et al.,^18^ which they had mutated to glutamine (here, H222Q-NOX4-Flag).

As shown in Fig. 4, when we subjected homogenates of cells expressing wt-NOX4-Flag to immunoprecipitation with anti-Flag antibodies, we brought down NOX4-Flag, and with it came abundant endogenous SOD3 protein. In contrast, anti-Flag immunoprecipitation of dd-NOX4-Flag or H222Q-NOX4-Flag from cell homogenates brought down mutant NOX4-Flag protein but with little or no SOD3 (Fig. 4). Thus, the two inserts in the extracellular E-loop of NOX4, and particularly histidine-222 in the first of those E-loop inserts, mediate the normal, robust binding of NOX4 to SOD3.

### Oxide hand-off from SOD3 to AQP3-PTEN

The surprising involvement of SOD3 resolves the first mystery from our introductory paragraphs – how NOX4 appears to generate H_2_O_2_ – but only magnifies the second – how can its product target PTEN? In other words, SOD3 is a cell-surface or extracellular enzyme (ecSOD); how does the H_2_O_2_ that it generates after insulin stimulation cross the plasma membrane, to enter the cell exactly where PTEN and PTPases reside, to quickly inactivate them without causing non-specific damage elsewhere? Contrary to the commonly held view that H_2_O_2_ is readily membrane-diffusible,^18,19,27^ the dipole moment (polarity) of H_2_O_2_ is slightly higher than water’s.^28^ This property makes non-facilitated diffusion of H_2_O_2_ through a hydrophobic lipid bilayer as slow or slower than non-facilitated diffusion of water.^28^ We inferred that there must be metabolic channeling of extracellular H_2_O_2_ from SOD3 through one or more transmembrane transporters, to target PTEN inside the cell.

A number of proteins have been implicated in the movement of reactive oxygen species across membranes (reviewed in reference^3^). Here, we eventually hypothesized a role for aquaporins (AQPs) in the oxide transport chain that controls balanced insulin signaling.^21^ To test our hypothesis, we began with low-dose AgNO_3_, a non-toxic, global inhibitor of facilitated diffusion through all aquaporins.^28,29^ In hepatocytes treated with AgNO_3_, we found that insulin failed to stimulate an intracellular burst of H_2_O_2_ (Extended Data Fig. 1a). Accordingly, insulin no longer caused the inactivation of PTEN inside these cells (Extended Data Fig. 1b). Insulin stimulation of AgNO_3_-treated hepatocytes provoked robust phosphorylations of ERK at Thr202 and Tyr204, as well as ample production of pT308-AKT, with only weak phosphorylation at the Ser473 site of AKT (Extended Data Fig. 1c). Extended Data Fig. 1d displays the effects of aquaporin blockade on targets downstream of AKT, i.e., continued insulin-stimulated phosphorylation of TSC2 (upstream of hepatic de-novo lipogenesis in Fig. 1), but poor insulin-stimulated phosphorylation of FOXO1 (required for suppression of gluconeogenesis). Thus, in the presence of a global inhibitor of diffusion through aquaporins, the red pathways in Fig. 1 remain insulin-responsive, while the blue pathways become insulin-resistant. This is the same pattern of imbalanced insulin action that occurs when we interfere with any other individual component of the oxide transport chain – namely, NOX4 (reference^6^) or SOD3 (Fig. 3). It is also the same deadly pattern that we saw in the liver in overnutrition and T2DM.^6^

### Mammals possess 13 aquaporins, numbered AQP0 through AQP12 (references^30,31^)

Here, we focused on the four aquaporins to date that have been reported to facilitate the diffusion of H_2_O_2_ across membranes – namely, AQP1, AQP3, AQP8, and AQP9. Literature on AQP1, AQP3, and AQP8 now appears conclusive that they each act as pores for H_2_O_2_ (references^3,28,32,33^). We also examined AQP9, owing to its abundance in hepatic parenchymal cells^31^ and its unusually promiscuous solute permeability that may^34^ or may not^28^ include H_2_O_2_·

**Figure 5:**
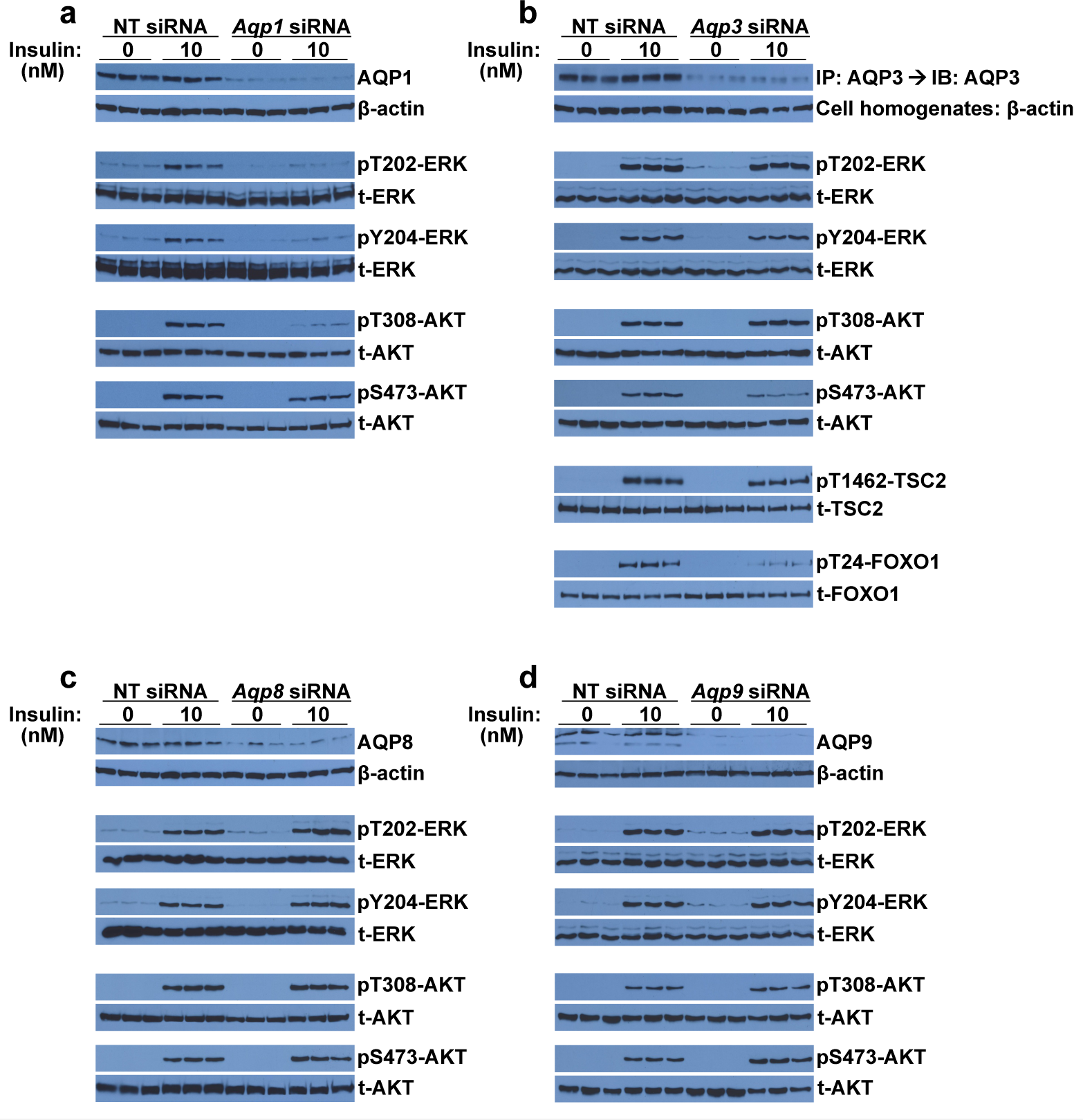
Solitary deficiency of AQP1 **(a)** or AQP3 **(b)**, but not of AQP8 (c) or AQP9 **(d)**, imbalances insulin signaling. As indicated, McArdle hepatocytes were pretreated with nontarget (NT) control siRNA or with siRNAs against individual aquaporins (*Aqp1*, *Aqp3, Aqp8*, or *Aqp9)*. Cells were exposed to 0 or 10 nmol/L insulin for 10 minutes and then harvested. For each aquaporin knock-down, the immunoblots displayed in this figure come from a single set of cultured cells. Shown are immunoblots of each of the four aquaporins to verify the knock-down, ß-actin to assess loading, insulin-stimulated phosphorylations of key sites on ERK (pT202-ERK, pY204-ERK) and AKT (pT308-AKT, pS473-AKT), and total ERK and AKT. In panel b, we also examined phosphorylated and total TSC2 and FOXO1. Each lane represents a separate culture well (n=3 per group).

Fig. 5a-d shows the effects of individual siRNA knock-downs of the four aquaporins of interest on insulin signaling to ERK, AKT, and selected AKT targets. Only the knockdown of AQP3 (Fig. 5b) recapitulated the distinctive pattern of imbalanced insulin signaling that we saw in the livers of hyperphagic T2DM animals^6^ and in cultured liver cells after knockdown or inhibition of NOX4 (reference^6^), knockdown of SOD3 (Fig. 3d), or global inhibition of facilitateddiffusion through all aquaporins (Extended Data Fig. 1c,d). As before, the pattern comprises two-site phosphorylation of ERK, the formation of pT308-AKT with impaired phosphorylation of Ser473 within AKT, continued insulin-stimulated phosphorylation of TSC2 (upstream of hepatic de-novo lipogenesis in Fig. 1), but poor insulin-stimulated phosphorylation of FOXO1 (required for suppression of gluconeogenesis).

Intriguingly, the knockdown of AQP1 (Fig. 5a) produced a mirror-image pattern of imbalanced insulin signaling – namely, strongly inhibited phosphorylations of both ERK sites and at Thr308-AKT, but fully responsive phosphorylation of Ser473 within AKT. This finding emphasizes that insulin-induced phosphorylation of AKT at Ser473 in liver cells behaves differently from phosphorylations of the two ERK sites and Thr308-AKT. Our knockdowns of AQP8 or AQP9 had no detectable effect on insulin signaling to either site on ERK or AKT (Fig. 5c,d).

Based on the functional data in Fig. 5b, we examined whether AQP3 physically associates with PTEN. Aquaporins are known to interact with a wide range of proteins,^35^ but an interaction with PTEN or PTPases had not been described. We exposed cultured McArdle hepatocytes to four conditions: pretreatment with non-target (NT) versus *Aqp3* siRNA, and then 10 minutes with 0 or 10 nM insulin. In these experiments, both AQP3 and PTEN were endogenous, i.e., at normal levels of expression in the cells pretreated with NT siRNA. To preserve the baseline structure of enzymatically active PTEN, which undergoes significant conformational changes upon oxidation and inactivation,^3,6,36–38^ we prepared cell extracts and then immunoprecipitated AQP3 under strictly anaerobic conditions within an enclosed workstation.

Immunoblots of this immunoprecipitated material show robust amounts of AQP3 and PTEN when prepared from cells with normal levels of AQP3 expression (NT siRNA) after exposure to either 0 (baseline) or 10 nM insulin (Fig. 6, leftmost six lanes). Normal baseline physical association of AQP3 with PTEN is consistent with our finding of a functional role for AQP3 in balanced insulin signaling (Fig. 5b). In other words, whenever insulin stimulates the NOX4-SOD3 complex to generate extracellular H_2_O_2_, AQP3 is already in position to channel this H_2_O_2_ across the cell membrane to PTEN. As verification of the specificity of the immunoprecipitation and immunoblotting antibodies that we used against AQP3, as well as the requirement for AQP3 protein to bring down PTEN, we found that signals for AQP3 and PTEN proteins were substantially attenuated in immunoprecipitates from cells pretreated with *Aqp3* siRNA (Fig. 6, rightmost six lanes). In contrast, knockdown of AQP3 did not alter the total amount of PTEN in whole-cell homogenates (Fig. 6).

**Figure 6:**
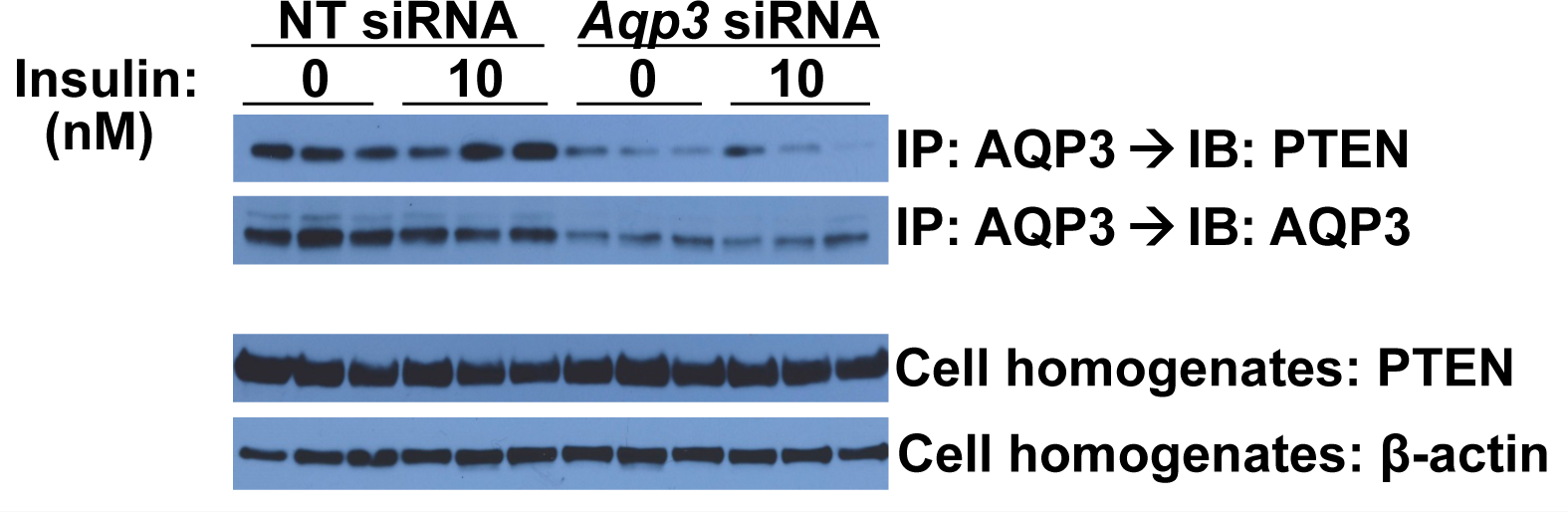
Aquaporin-3 forms a novel complex with PTEN. McArdle hepatocytes were pretreated with nontarget (NT) control or *Aqp3* siRNAs, as indicated, and then cells were exposed to 0 or 10 nmol/L insulin for 10 minutes. Cell homogenates were prepared and immunoprecipitation (IP) of AQP3 was performed under strictly anaerobic conditions within an enclosed workstation. Aquaporin-3 IP pellets were analyzed by immunoblotting (IB) for PTEN and, to verify the IPs and the knockdown, AQP3. To verify the starting material, whole-cell homogenates were analyzed by immunoblotting for PTEN and ß-actin (here) and for phosphorylated and total ERK and AKT (Extended Data Fig. 2). Displayed here are results from a single set of cultured cells. Each lane represents a separate well.

Insulin treatment had no effect on the association of AQP3 with PTEN (Fig. 6), indicating a stable complex during this period of insulin signaling, despite the large conformational changes of PTEN that occur upon its oxidation and inactivation. Extended Data Fig. 2 confirms that knockdown of AQP3 in this experiment caused the same harmful pattern of imbalanced insulin action that we saw in liver in overnutrition^6^ and in cultured hepatocytes after knockdown or inhibition of each component of the NSAPP oxide transport chain (NOX4 in reference^6^, SOD3 in Fig. 3d, AQPs globally in Extended Data Fig. 1c, and AQP3 in Fig. 5b).

## Discussion

Here, we report a new pathway of insulin signaling that we named the NSAPP oxide transport chain after its five major protein types: <u>N</u>OX4-<u>S</u>OD3-<u>A</u>QP3-<u>P</u>TEN/<u>P</u>TPases (schematic Extended Data Fig. 3). Thus, there are now <u>*three*</u> limbs of insulin signaling: the MAP kinase (ERK) limb, the metabolic (PI3K-AKT) limb, and the NSAPP oxide transport chain. We showed that the NSAPP pathway functions as a master regulator of normal, balanced insulin action via ERK, PI3K-AKT, and the downstream targets of AKT. Moreover, our work provides a molecular basis and precise functional role for the important but widely neglected “H_2_O_2_ hypothesis” of insulin signaling from the 1970s.^3,39^

Hepatic parenchymal cells and other cell types contain enzymes – PTEN and PTPases – that interfere with insulin signaling. Persistent activity of these enzymes was previously thought to globally block all downstream pathways, but instead, we reported crucial pathway-selective effects.^3,6^ Thus, for insulin to control hepatic glucose metabolism (blue in Fig. 1 and in Extended Data Fig. 3), the liver requires a robust system to inactivate PTEN and PTPases.^3,6^ The NSAPP oxide transport chain serves this function.

Our data show that impairment of any one of the components of the NSAPP oxide transport chain in cultured hepatocytes, from NOX4 up to PTEN, recapitulates all features of imbalanced insulin action that we^3,6,7,15^ and others^4,5,8–11,13,14^ have documented in states of overnutrition in vivo. These results imply that the molecular defect responsible for imbalanced insulin action in human T2DM liver^4^ may not necessarily reside in NOX4 itself, but instead could occur anywhere along the NSAPP oxide transport chain. In addition to the backbone proteins displayed schematically in Extended Data Fig. 3, the NSAPP oxide transport chain is also likely to interact with co-factors, such as proteins that restore PTEN activity after insulin signals have finished propagating (see section 4.5 in reference^3^). Key co-factors could also harbor molecular defects in obesity and T2DM.^3^

Not a single backbone member of the NSAPP oxide transport chain has shown up in any genome-wide association (GWA) study of obesity or T2DM (recently reviewed in reference^3^). Unfortunately, the current set of polymorphisms identified by GWA studies accounts for only about 10% of T2DM risk attributable to genetic factors, meaning significantly less than 10% of total disease risk, despite the high heritability of this disorder.^3,40^ Similarly, polymorphisms associated with obesity explain less than three percent of human variation in body-mass index.^41^ Thus, it is particularly important to seek novel molecular participants in obesity, the atherometabolic syndrome, and T2DM that have remained invisible to genome-wide surveys. Genes outside of the narrow set identified by GWA studies may explain the so-called missing heritability, i.e., why conditions such as obesity and T2DM that clearly exhibit high heritability in kindreds have proven largely intractable to GWAS.^3^

The importance of the NSAPP oxide transport chain is likely to extend beyond just insulin and its receptor. For example, other growth factor receptors, other tyrosine kinases, a number of G-protein coupled receptors, some membrane-bound oncogenes, and the long form of the leptin receptor (LEPRb) have been shown to signal intracellularly via the phosphorylation of specific tyrosyl residues and/or the generation of specific 3’-phosphoinositides.^42–44^ For these pathways to fully propagate, they would seem to require highly regulated inactivation of nearby PTEN and PTPases that would otherwise digest these specific phosphorylated sites. In fact, normal signaling by several growth factors and by leptin has been reported to depend on uncharacterized oxidants or even on H_2_O_2_ itself.^3,24,32,45–49^

Our results imply that the activities of NOX4, SOD3, and AQP3 are required for normal insulin signaling to Ser473-AKT. The role of AQP1 appears to be more structural: its physical presence in liver cells is required for insulin signaling to ERK and to Thr308-AKT, but global inhibition of transmembrane diffusion through all aquaporins by low-dose AgNO_3_ left signaling to those targets fully intact. Based on these results, we infer that AQP1 may serve additional functions beyond acting as a pore, e.g., possibly as a scaffold.

In conclusion, the NSAPP oxide transport chain serves essential functions in balanced hormonal signaling. Now that we know of the existence of the NSAPP pathway, unraveling the molecular basis for its dysfunction in overnutrition, obesity, the atherometabolic syndrome, and T2DM has become a top priority.

## End notes

**Supplementary information** is available in the submitted version of this manuscript.

## Acknowledgements

The authors acknowledge support from a recruitment package from the Temple University School of Medicine in 2009, Basic Science Award #1–13-BS-209 from the American Diabetes Association (2013–2015), the Ruth and Yonatan Ben-Avraham Fund, and the Swedish Heart-Lung Foundation (Hjärt-Lungfonden). We are particularly grateful to these organizations, because the National Institutes of Health (NIH-USA) have repeatedly refused to support our work on the molecular basis for pathway-selective insulin resistance and responsiveness. This last point is consistent with the well-documented practice of the NIH to reward scientific conformity.^50,51^

## Author contributions

X.W. optimized all methods and performed all experiments except those in Fig. 3. K.C. optimized the methods and performed the experiments in Fig. 3. K.J.W. secured funding, designed the experiments, reviewed all primary data, drew the schematics in Fig. 1 and in Extended Data Fig. 3, and wrote the manuscript.

## Author information

K.J.W. reports an ownership interest in Hygieia, Inc., and in Gemphire Therapeutics, Inc., and serves on the Medical and Scientific Advisory Board of Gemphire Therapeutics, Inc. The other authors declare no competing financial interests. Correspondence and requests for materials should be addressed to K.J.W.

## Methods

### Reagents

McArdle 7777 rat hepatoma cells were obtained from the American Type Culture Collection (Manassas, VA, USA; catalog no. CRL-1601) and cultured as described previously.^6,52,53^ We had chosen these cells, because they respond to a physiologically plausible concentration of insulin, 10nmol/L, by robustly phosphorylating every downstream phosphorylation target of the insulin receptor that we examined.^6^ Moreover, we have found these cells to be readily transfected with siRNAs and expression plasmids.^6,52–54^

Antibodies against target proteins (total target as well as forms with site-specific phosphorylations) are listed in Extended Data Table 1, following the nomenclature in Fig. 1 and Extended Data Fig. 3 (the latter drawn to include a schematic representation of the NSAPP oxide transport chain). To achieve reproducible, high-quality immunoprecipitations (IP) and immunoblots (IB) of the proteins in the NSAPP pathway, we had to test over 50 commercially available antibodies against NOX4, SODs, AQPs, and PTEN, most of which failed to work properly. This vexing problem has been discussed at length elsewhere.^55,56^ Hence, in Extended Data Table 1, we list the target protein, the epitope if known, the supplier and catalogue number of the antibody that we successfully used, a description of the antibody including its clonality, our use of it (IP or IB), and the molecular weight of the target. Additional information can be found in the methods section and first supplemental table of our initial article on NOX4 and SEIRR.^6^ The antibody against NOX4 that we used in that earlier study^6^ is no longer available, and so we had to find a new one.

The antibody set for aquaporins-1, -3, -8, and -9 required additional attention. Using siRNAs, described below, we were able to validate a set of antibodies that detect each of these AQPs in immunoblots, based on known molecular weights and fading of the signal after optimized use of specific siRNAs (Figs. 5 and 6). Detection of AQP3 in McArdle cell homogenates required prior immunoprecipitation of this protein, an approach that we and others have previously used to improve sensitivity and specificity of immunoblots.^6^

### Molecular and chemical manipulations of the NSAPP oxide transport chain in cultured liver cells

Following our previously published knock-down protocol,^6^ McArdle hepatoma cells replete or deficient in individual components of the NSAPP oxide transport chain were prepared by transfection with nontarget siRNA (designed and microarray tested by the manufacturer for minimal targeting of rat genes) or siRNAs against the rat target of interest (SmartPool, Dharmacon, Lafayette, CO, USA), using the siIMPORTER transfection reagent (Millipore Corporation, Billerica, MA, cat # 64–101). With one exception, all siRNAs were used at a concentration of 50 nmol/L for 4 hours in serum-free medium and then overnight in DMEM with 10% FBS, followed by an additional 48 hours at 37°C in DMEM/10% FBS without siRNA (three days total), as previously described.^6^ Successful knock-down of AQP8, however, required target and nontarget siRNAs at 100nmol/L and then 72, not just 48, hours at 37°C in DMEM/10% FBS without siRNA (four days total). Catalogue numbers and targeted mRNA sequences are listed for each siRNA in Extended Data Table 2.

Cells were approximately 40% confluent when the siRNAs were added, so that confluence would be approximately 80% during the insulin signaling studies three or four days later. Cells were switched to serum-free medium (DMEM/1% BSA) for 3 hours before supplementation with 0 or 10 nmol insulin/L. Exposure to 0 or 10 nmol insulin/L lasted 10 minutes, to allow intracellular H_2_O_2_ accumulation, PTEN inactivation, and phosphorylations of preexisting protein targets. To chemically block aquaporin channels, we used a low concentration, 10μmol/L, of a non-toxic global aquaporin inhibitor, AgNO_3_ (references^28,29^), which we added to cells 15 min before supplementation with insulin. The AgNO_3_ remained on the cells during the subsequent 10 min of insulin exposure.

### Assessments of key targets within each limb of insulin signaling, i.e., the ERK and AKT limbs and the NSAPP oxide transport chain

To assess insulin-stimulated accumulation of intracellular H_2_O_2_, McArdle hepatocytes were plated in collagen-coated, glass-bottom Petrie dishes (MatTek Corp. Ashland, MA, cat # P35GCOL-0–14-C) designed for use in confocal microscopy. Cells were pre-treated with siRNAs or AgNO_3_, as described above, given 0 or 10 nmol insulin/L, and then immediately supplemented with CellROX Deep Red (5μM final concentration, Invitrogen, Carlsbad, CA, cat # C10422), an intravital fluorescent dye that detects a variety of oxides. CellROX is cell-permeable and non-fluorescent until it reacts with intracellular oxides, at which point it becomes cell-impermeable and fluorescent. After 10 min, the cells were washed, immediately fixed in 3.7% formaldehyde for 15min according to the manufacturer’s instructions, and imaged within two hours on a confocal microscope (Leica Microsystems LAS AF, Buffalo Grove, IL), using a plane selected at the level of the cell nuclei. Prompt confocal microscopy of fixed samples produces high-resolution images. To assess the component of the CellROX signal that is specifically attributable to H_2_O_2_, the media in some of the Petrie dishes were supplemented with native bovine liver catalase (final concentration 8 mg/ml, Sigma, St. Louis, MO, cat # C-1345), a highly specific enzyme, two minutes before the addition of insulin and CellROX Deep Red. The exogenous catalase remained on the cells during the subsequent 10 min of insulin exposure. Native catalase has the additional advantage that, when added to culture medium, the enzyme shows no or nearly no cellular adherence or entry^57^ and can therefore test extracellular exposure of H_2_O_2_ in the NSAPP oxide transport chain (Extended Data Fig. 3 and reference^3^).

Inactivation of PTEN was assessed as we previously described, through the use of immunoprecipitation and then enzymatic assays within a strictly anaerobic workstation to preserve the true activity.^3,6,37,46^ Phosphorylations of each insulin-responsive site on ERK (T202, Y204) and AKT (T308, S473) were assessed by immunoblotting, using site-specific antibodies, as we described previously.^6^ Insulin-stimulated phosphorylations of targets directly downstream of AKT (total target as well as forms with site-specific phosphorylations) were also assessed by immunoblotting (see Fig. 1, Extended Data Table 1, and reference^6^). All membranes used for immunoblots of phosphorylated proteins were stripped and reprobed for the total target protein. In some cases, the immunoblot image after stripping was of poor quality, and so we prepared separate gels and membranes using the same amount of the same cellular homogenates. Each insulin signaling experiment was performed at least three independent times.

To assess NSAPP structure, we performed co-immunoprecipitations following our published protocols.^6,52,53^ To ensure that we used similar amounts of starting material, we quantified total protein concentrations in whole-cell homogenates and then used the same amounts of whole-cell protein for immunoprecipitations. We also performed immunoblots of these same whole-cell homogenates to detect ß-actin. For co-immunoprecipitations that gave negative results, we included lanes of whole-cell extracts on the same immunoblot, to verify that the molecule that did not measurably co-immunoprecipitate was, in fact, present in the original samples at levels detectable by our antibodies.

### Expression vectors for Flag-tagged NOX4 and site-directed mutants of NOX4-Flag

A plasmid containing cDNA that encodes the full-length wild-type (wt) human NOX4 protein was purchased from Addgene (Cambridge, MA, USA; plasmid # 69352, pcDNA3.1-hNox4, originally deposited by the Krause laboratory^17^). We used high-fidelity PCR to amplify the *NOX4* cDNA while inserting DNA encoding the Flag epitope tag (dykddddk)^58^ just before the stop codon. In the same PCR, we also added sites for restriction enzymes after the stop codon (our PCR primer sequences are designated NOX4+Flag and given in Extended Data Table 3). Digestion of the amplimer and the expression vector pcDNA3.1(-) (Invitrogen/Thermo-Fisher) was followed by ligation, bacterial transformation, selection of a single clone, and verification of the sequence of the wt-NOX4-Flag insert by a commercial service (Genewiz, South Plainfield, NJ, USA).

Next, we created a double-deletion mutant (dd-NOX4-Flag) that lacks both of the E-loop inserts that had been identified by Takac et al.^18^ We used the QuikChange kit (catalog no. 200518, Stratagene-Agilent Technology, Santa Clara, CA),^53,54^ with our wt-NOX4-Flag expression plasmid as the starting template and mutagenesis primers that we had optimized after several attempts (these primers are also listed in Extended Data Table 3). The longer E-loop insert at the 5’ end (18 codons) was removed first (primers designated d-E-loop-1), and then the process was repeated to remove the shorter E-loop insert as well (10 codons; primers d-E-loop-2). We also made a site-directed mutant of the key histidine-222 residue in the first E-loop insert to glutamine (primers H222Q, to make the H222Q-NOX4-Flag construct; see Extended Data Table 3).^18^ All mutants were sequenced to verify the introduction of DNA changes.

The empty pcDNA3.1(-) vector, the unmutated wt-NOX4-Flag plasmid, and the two mutant plasmids (dd-NOX4-Flag and H222Q-NOX4-Flag) were transfected into separate cultures of McArdle cells, using the FuGENE 6 reagent (Promega Corporation, Madison, WI, USA, cat # E2691). Stably expressing McArdle clones were selected with G418, followed by verification of protein expression by immunoblots of whole-cell homogenates using anti-Flag antibodies to detect bands of the appropriate molecular weights. We chose cell lines with similar levels of expression, based on anti-Flag immunoblots of whole-cell homogenates.

### Statistics

Normally distributed data are reported as means±SEMs. For comparisons involving several groups of cultured hepatocytes, ANOVA was initially used, followed by pairwise comparisons using Tukey’s test (Prism 7, GraphPad Software, California, USA).

## Extended data

**Extended Data Figure 1:**
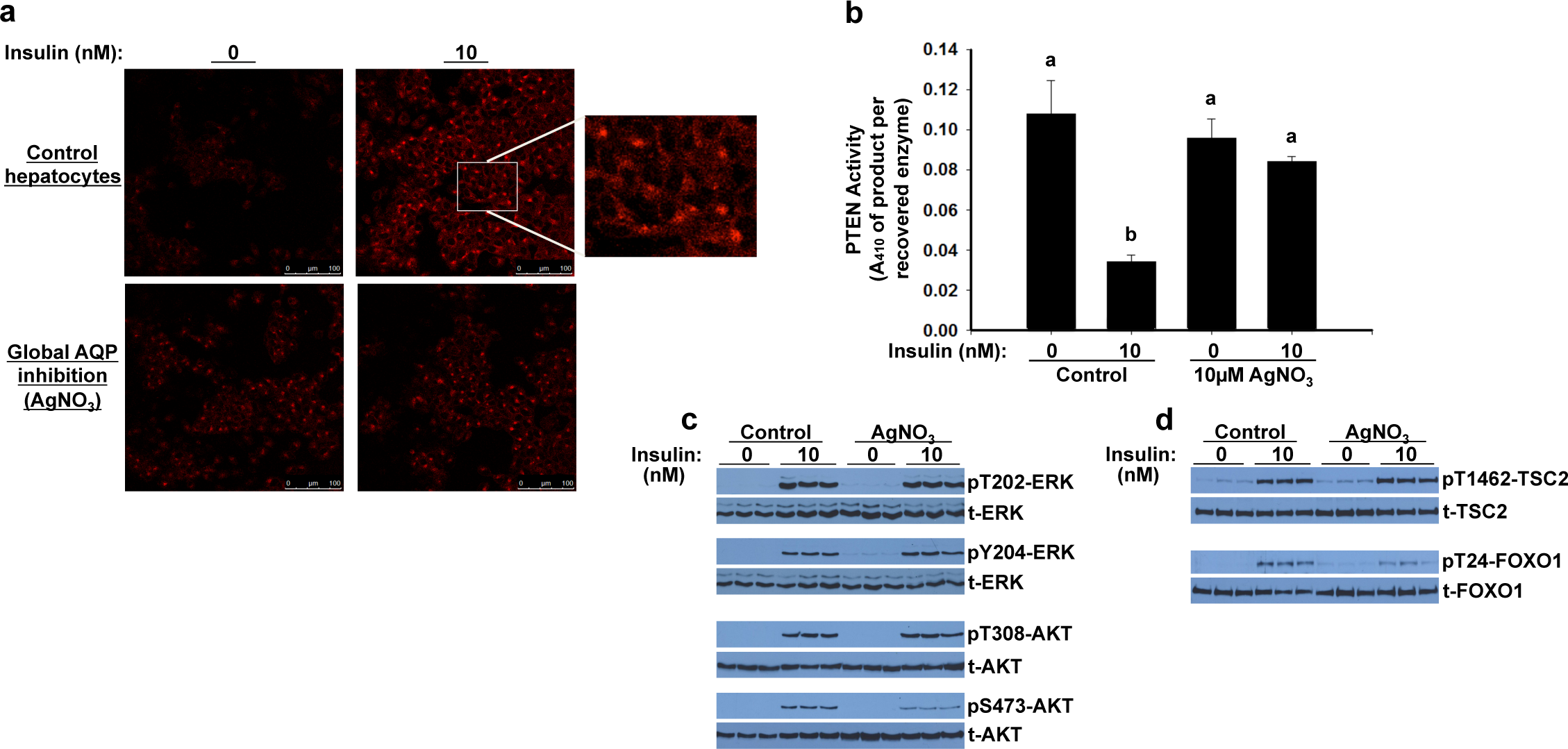
Global inhibition of facilitated diffusion through aquaporins in cultured McArdle hepatocytes recapitulates the harmful pattern of imbalanced insulin signaling seen in overnutrition or after artificial disruption of NOX4 or SOD3. As indicated, cells were pre-treated with 0 (Control) or 10μM AgNO_3_, a non-toxic, global inhibitor of facilitated diffusion through all aquaporins, and then exposed to 0 or 10 nmol/L insulin for 10 minutes and harvested. Immunoblots came from a single set of cultured cells. Each lane represents a separate well. **a**, Insulin-stimulated production of intracellular H_2_O_2_, assessed by confocal images of CellROX Deep Red fluorescence. We definitively identified the intracellular CellROX signal as H_2_O_2_ by its quenching by catalase (not shown). As in Fig. 3b, the insulin-stimulated burst of intracellular H_2_O_2_ in individual control hepatocytes was localized mainly to a perinuclear area (inset). **b**, PTEN activities in cell homogenates, assayed under strictly anaerobic conditions (means±SEMs, n=3, P=0.0034 by ANOVA; columns that do not share a lowercase letter differ by Tukey’s test, P<0.05). **c**, Insulin-stimulated phosphorylations of key sites on ERK and AKT. Total (t-) ERK, AKT are also shown. **d**, Insulin-stimulated phosphorylations of targets directly downstream of AKT, in de-novo lipogenic pathways (pT1462-TSC2) and in glucose-lowering (pT24-FOXO1). Nomenclature follows Fig. 1.

**Extended Data Figure 2:**
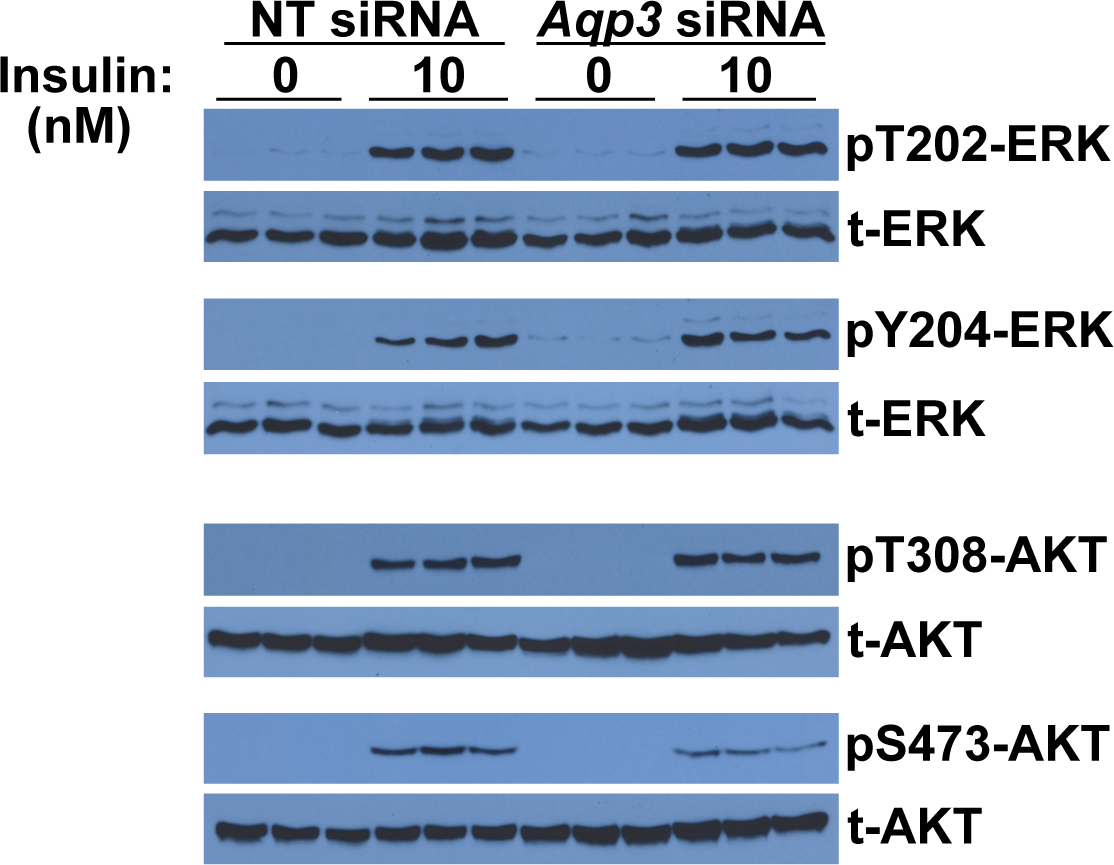
Knock-down of AQP3 in cultured hepatocytes recapitulates the harmful pattern of imbalanced insulin signaling seen in overnutrition in vivo or after artificial disruption of NOX4 or SOD3 or global inhibition of aguaporins in vitro. Cell homogenates used in Fig. 6 were analyzed by immunoblotting for insulin-stimulated phosphorylations of key sites on ERK (pT202-ERK, pY204-ERK) and AKT (pT308-AKT, pS473-AKT), and total ERK and AKT. Each lane represents a separate culture well, aligned with Fig. 6 (n=3 per group). These data confirm that the signaling defect that we found in the AQP3-deficient cells of Fig. 5b also occurred in the AQP3-deficient cells of Fig. 6.

**Extended Data Figure 3:**
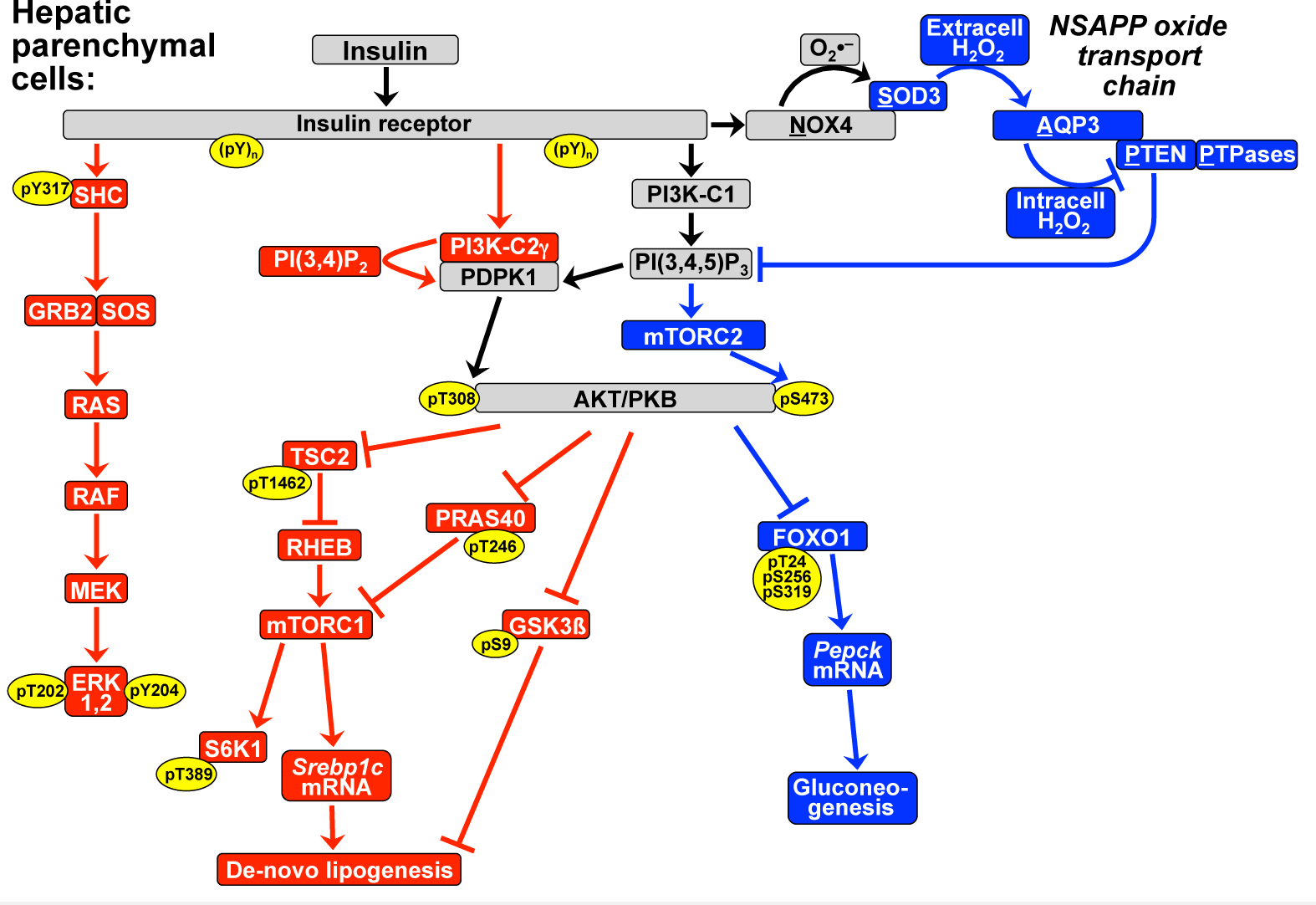
New model of the three key limbs of insulin signaling in hepatic parenchymal cells. Here we summarize the findings of the current study by replacing the encircled pathway from NOX4 to PTEN in Fig. 1 with a schematic of the NSAPP oxide transport chain. The NSAPP pathway is shown in context with the other key limbs of insulin signaling in hepatic parenchymal cells. The insulin receptor, NADPH oxidase-4 (NOX4), and aquaporin-3 (AQP3) are transmembrane proteins. Everything shown above them is extracellular, and everything shown below them is intracellular. Extracell, extracellular; Intracell, intracellular; 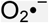, the superoxide anion; SOD3, superoxide dismutase-3. Other nomenclature follows Fig. 1. Adapted from references^3,6^ with permission.

**Extended Data Table 1:**
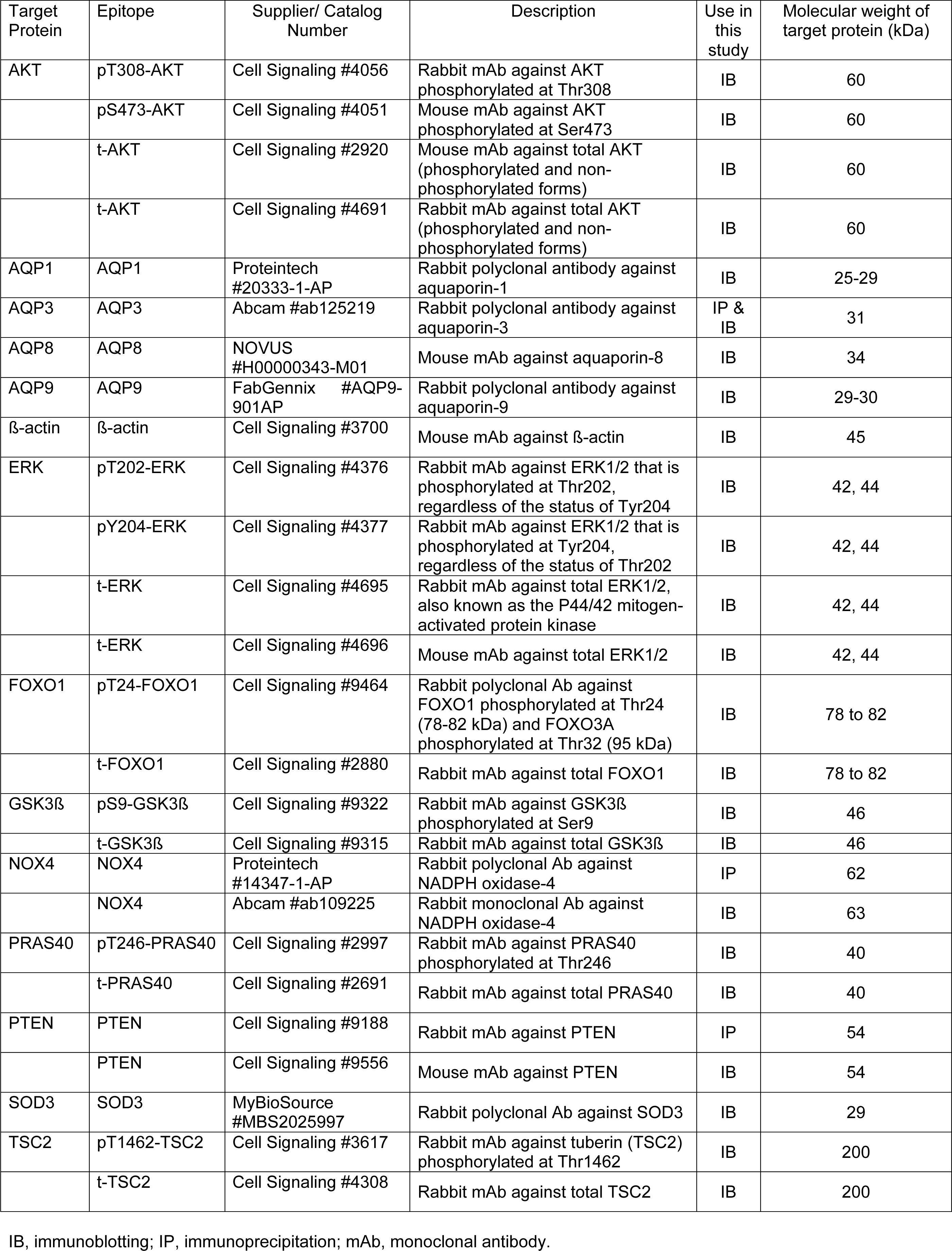
Antibodies against target proteins and epitopes.

**Extended Data Table 2:**
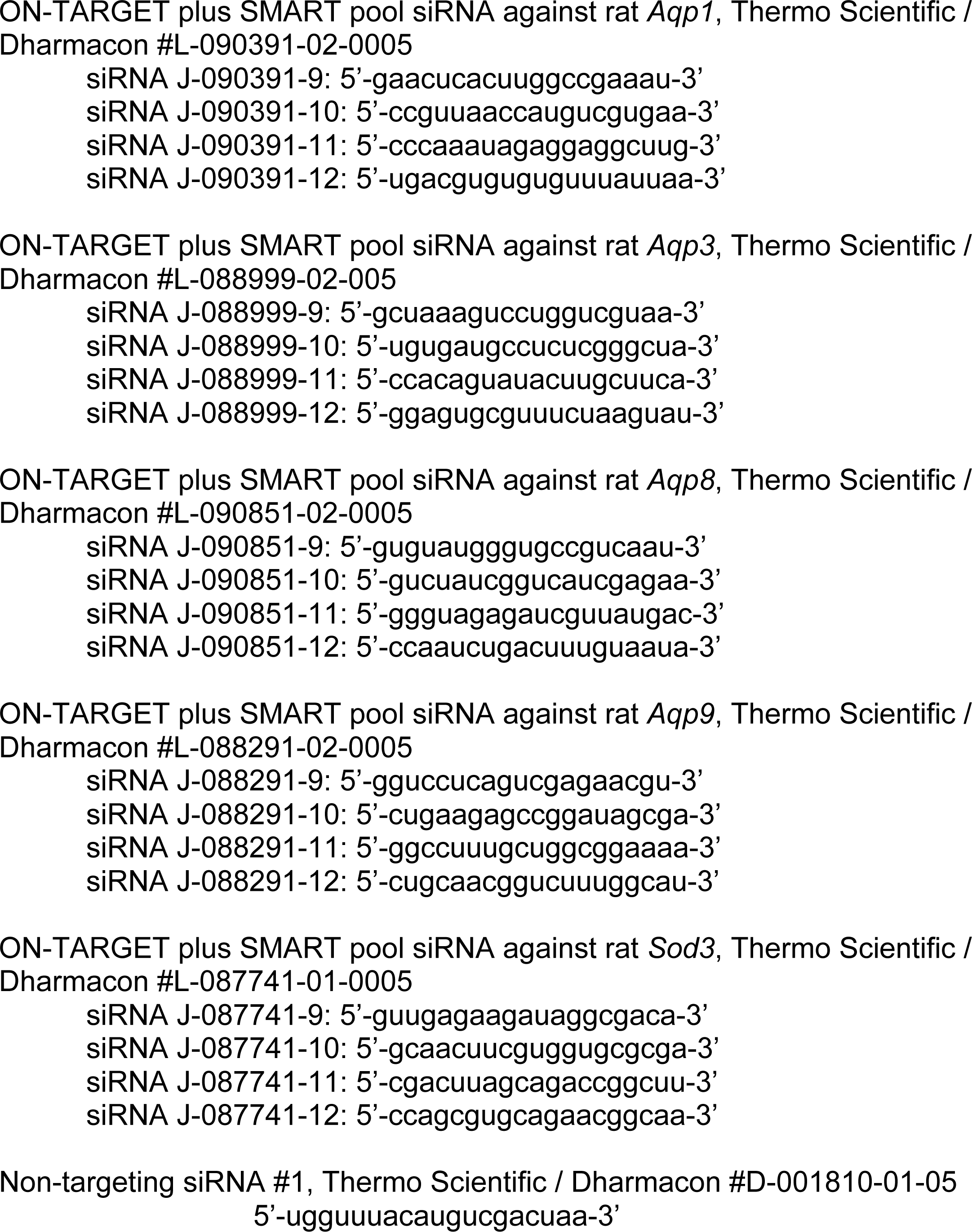
Messenger RNA sequences targeted by small interfering (si) RNAs.

**Extended Data Table 3:** Optimized primer sequences for PCR amplification and site-directed mutagenesis of human *NOX4* cDNA.

| Primer names<br>(from Methods) | Forward primer | Reverse primer |
| --- | --- | --- |
| NOX4+Flag | 5'-aataataat <u>ctcgag</u><br>gggccc **atg** gct gtg tcc<br>tgg agg agc tgg ctc gcc<br>aac-3' | 5'-ggcggcggc <u>ggtacc tta</u><br><u>ctt gtc atc gtc gtc ctt gta</u><br><u>gtc gct gaa aga ctc ttt att</u><br>gta ttc aaa tc-3' |
| d-E-loop-1 | 5'-ggg ctg ctg aag tat caa<br>act—aat att tcc tta cca<br>gag tat ttc-3' | 5'-gaa ata ctc tgg taa gga<br>aat att—agt ttg ata ctt cag<br>cag ccc-3' |
| d-E-loop-2 | 5'-ca aaa ccg gca gag ttt<br>acc cag—ccc aga ttc caa<br>gct aat ttt c-3' | 5'-g aaa att agc ttg gaa tct<br>ggg—ctg ggt aaa ctc tgc<br>cgg ttt tg-3' |
| H222Q | 5'-caa act aat tta gat acc<br>**cag** cct ccc ggc tgc atc<br>agt c-3' | 5'-g act gat gca gcc ggg<br>agg **ctg** ggt atc taa att agt<br>ttg-3' |

Codons are set off from each other; restriction enzyme sites and the complement of the sequence encoding the Flag epitope tag are underlined; start, stop, and mutated codons or anti-codons are bolded; and the location of each site-directed deletion is indicated by an em dash.

